# Loss of H3.1K27me1 in Arabidopsis confers resistance to Geminivirus by sequestering DNA repair proteins onto rDNA and defense-related genes

**DOI:** 10.1101/2022.09.13.507805

**Authors:** Zhen Wang, Claudia M. Castillo Gonzalez, Changjiang Zhao, Chun-Yip Tong, Changhao Li, Zhiyang Liu, Kaili Xie, Jiaying Zhu, Zhongshou Wu, Xu Peng, Yannick Jacob, Scott D. Michaels, Steven E. Jacobsen, Xiuren Zhang

**Affiliations:** Department of Biochemistry and Biophysics, Texas A&M University, College Station, TX 77843, USA.; Molecular and Environmental Plant Sciences, Texas A&M University, College Station, TX 77843, USA.; Department of Molecular, Cell, and Developmental Biology, University of California, Los Angeles, Los Angeles, CA 90095, USA; Howard Hughes Medical Institute, University of California, Los Angeles, Los Angeles, CA 90095, USA; Department of Molecular Physiology, College of Medicine, Texas A&M University, College Station, TX 77843, USA; Department of Molecular, Cellular & Developmental Biology, Yale University, New Haven, CT, 06511; Department of Biology, Indiana University, Bloomington, IN 47405; Department of Biology, Texas A&M University, College Station, TX 77843, USA

**Keywords:** Histone methyltransferase, H3.1K27me1, Genomic Stability, Transcription Associated Homologous Recombination Repair, RAD51, RPA1A, Geminivirus

## Abstract

The H3 methyltransferases ATXR5 and ATXR6 deposit H3.1K27me1 to heterochromatin to prevent genomic instability and transposon reactivation. Here, we report that *atxr5 atxr6* mutants displayed robust resistance to Geminivirus. The viral resistance correlated with activation of DNA repair pathways, but not with transposon reactivation or heterochromatin amplification. We identified RAD51 and RPA1A as partners of virus-encoded Rep protein. The two DNA repair proteins showed increased binding to heterochromatic regions and defense-related genes in *atxr5 atxr6* vs wild type plants. Consequently, the proteins had reduced interactions to viral DNA in the mutant, thus hampering viral replication. Additionally, RAD51 recruitment to the host genome arose via BRCA1, HOP2 and CYCB1, and this recruitment was essential for viral resistance in *atxr5 atxr6*. Thus, Geminiviruses adapt to healthy plants by hijacking its DNA repairing pathways for replication, but the host could retain DNA repairing proteins via sacrificing its genome stability to suppress viral infection.

## Introduction

Geminiviruses, a group of single stranded circular DNA viruses, has been considered an increasing threat to crop yield and food security worldwide due to their broad host range^1, 2^. Epigenetic modifications, including histone methylation, regulates various development processes and plant defense responses against microbes such as plant viruses^3, 4^. For example, kryptonite (KYP)/SUVH4 catalyzes the deposition of H3K9me2/3 on viral mini-chromosome to attenuate virus accumulation, whereas Geminivirus-encoded transcriptional activation protein (TrAP) can impair the KYP activity to counter the host defense ^3^. Recently, various histone marks such as H4K3me3, H3K36me3 and H3K27me3 has been detected on Geminivirus minichromosomes^5, 6^, suggestive of regulatory roles for histone methylation in plant defense against Geminiviruses.

Arabidopsis Trithorax-Related Proteins 5 (ATXR5) and ATXR6 (ATXR5/6) redundantly catalyze K27 monomethylation specifically on the H3.1 variant (H3.1K27me1) genome-wide, but depletion of this mark impact mainly heterochromatin^7–9^. Lower levels of H3.1K27me1 in heterochromatic regions cause increased amount of H3.1K27ac and H3.1K36ac, leading to transcriptional activation of transposable elements (TEs)^8, 10^. Reduction of H3.1K27me1 also generates widespread DNA amplification in heterochromatic regions of the genome^8, 9^. DNA duplication in *atxr5 atxr6* is associated with the accumulation of DNA double strand breaks (DSBs)^7, 8, 11, 12^. DSBs can be repaired through homology recombination repair (HRR) and non-homologous end joining (NHEJ) in eukaryotic cells. It has been found that HRR factors such as Arabidopsis *RAD51*, breast cancer susceptibility gene 1 (*BRCA1*), and *CYCB1* are transcriptionally upregulated in *atxr5 atxr6* mutants^11–13^. Intriguingly, a genetic screen based on the readout of *RAD51* promoter-driven GFP in the *atxr5 atxr6* background recovered loss of function mutants of *METHYL-CpG BINDING DOMAIN PROTEIN 9* (*MBD9*) and *Yeast SAC3 HOMOLOG B (SAC3B)* as suppressors of *atxr5 atxr6* phenotypes. These lines provide an opportunity to investigate the mechanisms underlying the distinct molecular phenotypes of *atxr5 atxr6*^12, 14^.

Recent work has revealed that H3.1 interacts with TONSOKU (TSK), which is required for initiating HRR during replication to resolve stalled/broken replication forks and maintain genomic stability^15^. Interestingly, inactivation of TSK in *atxr5 atxr6* mutants suppresses heterochromatin amplification, whereas deletions of RAD51 and BRCA1 enhance this phenotype^12, 15^. These results suggest that the roles of different HRR proteins in protecting genome stability may be distinct. Furthermore, there is a still a gap in our understanding of the interplay between H3.1K27me1 depletion, heterochromatin amplification and HRR.

A growing body of evidence highlights the involvement of HRR factors in viral DNA replication in human^16–21^. In plants, roles of HRR factors in geminiviral propagation are perceived in different ways. Whereas proliferating cell nuclear antigen (PCNA) suppresses the enzyme activity of Rep, RAD54 promotes Rep function in vitro. Moreover, deficiency of PCNA or RAD51D impairs Geminivirus accumulation, but deletion of RAD54 or RAD17 does not affect infection^22–26^. The contrasting reports above indicate that the roles of plant HRR factors in viral DNA replication remain unclear. Of note, when the plant innate immune response is triggered by salicylic acid (SA), RAD51 can directly bind to promoter elements of defense genes and enhance gene expression in a *BRCA2-* and SA-dependent manner. Moreover, *RAD17* and *Rad-3-related* (*ATR*) are required to enhance the expression of SA-activated genes and deploy an effective immune response^27, 28^. It has been reported that Cabbage Leaf Curl Virus (CaLCuV) infection can induce the expression of genes in the SA pathway^29^, which in turn seesaws the battle between Geminiviruses and plants^30, 31^. These results suggest that HRR proteins might regulate the plant defense against Geminiviruses through the innate immune response.

We have recently surveyed the viral susceptibility of numerous epigenetic mutants of Arabidopsis. Surprisingly, we found that *atxr5 atxr6* double mutants behaved differently and displayed a striking resistance to CaLCuV. Depletion of *SAC3B*, *MBD9* or *BRCA1* in *atxr5 atxr6* restored susceptibility of the viral infection despite contrasting effects on TE reactivation and heterochromatin amplification. Transcriptome-wide association studies (TWAS) showed that reduced viral DNA replication correlated with upregulation of HRR related genes in the mutants. We found that the viral protein Rep hijacked host RAD51 and RPA1A on the viral genome to promote viral amplification. Interestingly, RAD51 and RPA1A showed robust binding to unstable genomic DNA (e.g., rDNA and noncoding RNA (ncRNA) loci) in *atxr5 atxr6*. Moreover, RAD51 was enriched at plant defense genes, and its binding was coupled with transcriptional upregulation in *atxr5 atxr6*. Additionally, we found that BRCA1, HOP2 and CYCB1 recruit RAD51 onto the host genome and deletion of these factors restored the susceptibility of *atxr5 atxr6* to geminiviral infection. Thus, we propose that unstable genomic DNA exemplified by rDNA, together with defense-related gene loci, sequesters RAD51 via BRCA1, HOP2 and CYCB1 to prevent loading of HRR factors onto viral genome, leading to poor viral replication.

## Results

### The *atxr5 atxr6* mutant displays increased resistance to CaLCuV infection

To investigate the effect of epigenetic modifications on viral pathogenesis, we inoculated numerous mutants in epigenetic silencing pathways of Arabidopsis with CaLCuV. The symptoms of CaLCuV infection included chlorosis, curled leaf and plant growth arrest (Fig. S1A). Loss-of-function mutants of H3K9me2/3 methyltransferases (MTases) (*suvh4/5/6*), H3K27me3 MTases in Polycomb Repressive Complex 2 (PRC2) (*clf-28*) and DNA MTases (*drm1 drm2 cmt3*) displayed increased viral susceptibility compared to wild type (WT) Col-0, implying their critical roles in inhibiting viral propagations in plants (Extended Data Fig. 1). In contrast, assays in *atxr5 atxr6* mutants (with compromised H3.1K27me1 deposition) resulted in significantly fewer infected plants, and plants displaying milder symptoms and lower viral titers compared to either of the single mutants of *atxr5* and *atxr6* or Col-0 (Fig. 1a). These results indicated that *atxr5 atxr6* did not offer a permissible environment for Geminivirus infection and/or replication (Fig. 1a, b).

**Fig. 1.**
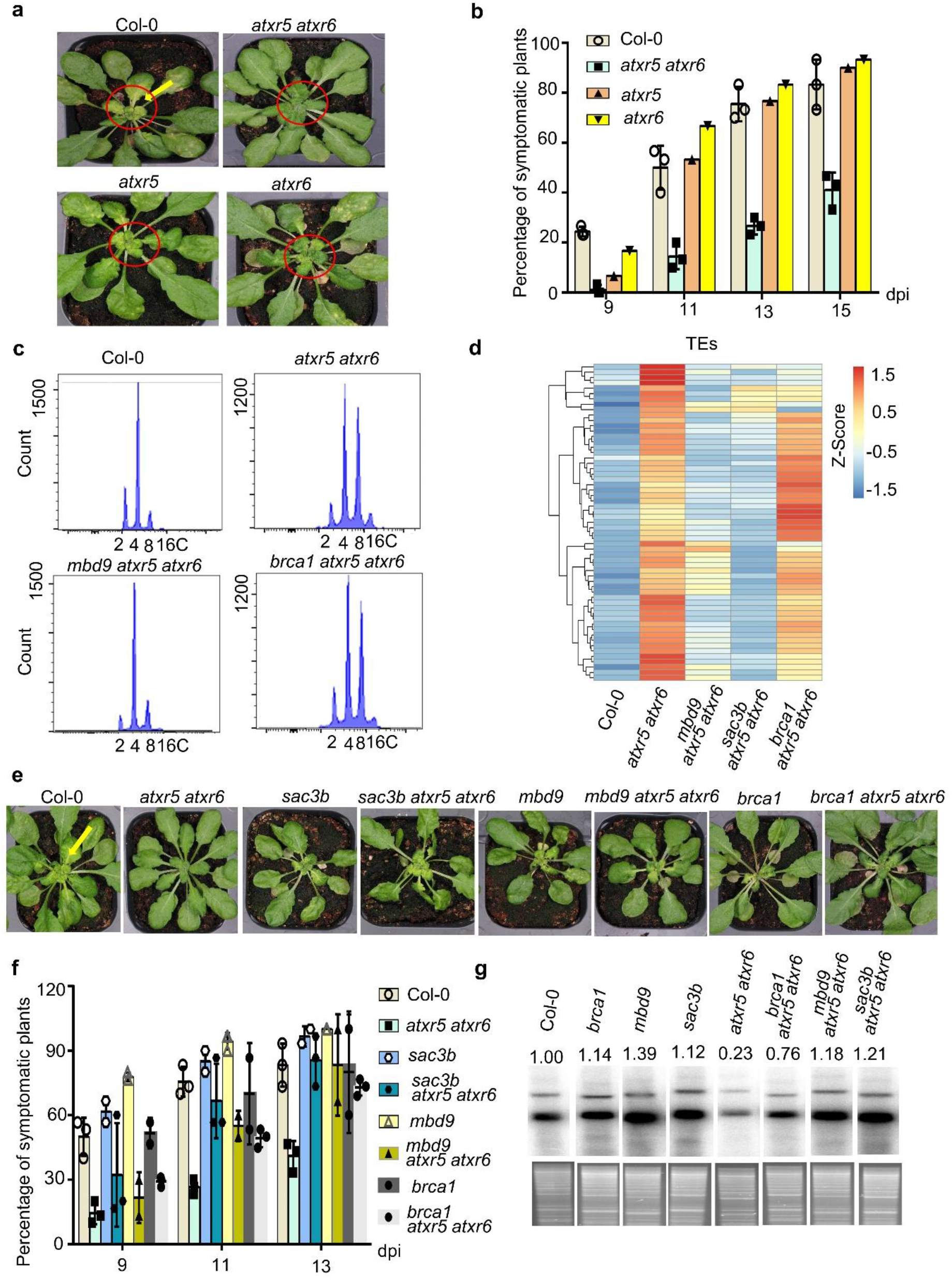
*atxr5 atxr6* displays robust viral resistance phenotype that is superficially uncoupled with re-activation of TEs and DNA replication. **a,** Loss-of-function mutants of *atxr5 atxr6* show resistance to CaLCuV. Photographs of CaLCuV-infected Col-0, *atxr5*, *atxr6* and atxr5 atxr6 plants at 16 dpi. **b,** Percentages of symptomatic plant induced by CaLCuV-infection at 9, 11, 13 and 15 dpi. Each dot in the bar plot represents one replicate, experiments were performed with 36 plants/replicate. **c,** Flow cytometry assays shows that loss of MBD9 but not BRCA1 could rescue DNA re-replication phenotype in *atxr5 atxr6*. **d,** Heat map shows that loss of MBD9, or SAC3B, but not BRCA1, could suppress transcriptional re-activation of TEs in *atxr5 atxr6*. The quantification was conducted by de-seq2. **e,** The loss of MBP9, SAC3B, or BRCA1 could all increased viral susceptibility of *atxr5 atxr6*. Photographs of CaLCuV infected plants were taken at 14 dpi. **f,** Percentages of symptomatic plants in different backgrounds induced by CaLCuV-infection at 9, 11 and 13 dpi. Each dot in the bar plot represents one replicate, experiments were performed with 36 plants/replicate. **g,** Southern blot assays showed that the accumulation of viral DNA A in CalCuV-infected plants with different genotypes at 14 dpi. EcoRI-digested input DNA serves as a loading control (Bottom panel).

As the infection was conducted through agrobacteria-mediated infiltration, we first examined whether the initial plasmid delivery to planta was compromised in *atxr5 atxr6*. We collected inoculated leaves at 3, 6, 9 and 13 dpi (day post inoculation). Southern blot, semi-qPCR and qPCR results showed that the amount of delivered plasmids in Col-0 and *atxr5 atxr6* was comparable at 3, 6, 9 and 13 dpi. In contrast, the amount of replicated viral DNA was strikingly lower in *atxr5 atxr6* vs. Col-0 starting at 13 dpi (Extended Data Fig. 2). These results indicated that viral DNA replication rather than plasmid transfection was suppressed in *atxr5 atxr6*.

ATXR5/6 deposit K27me1 specifically on the replication-dependent H3.1 variant, which prevents heterochromatin amplification^9^. H3.1 is encoded by five genes in *Arabidopsis thaliana,* and inactivation of H3.1 in plants leads to sterility and strong pleotropic phenotypes^32^. Deletion of *FASCIATA2* (*FAS2*), which encodes a subunit of CHROMATIN ASSEMBLY FACTOR 1 (CAF1), prevents the normal deposition of H3.1 during replication^9, 33^. Of note, while *fas2* and *h3.1* mutants lose both H3.1K27me1 and the H3.1 variant, the *atxr5 atxr6* mutant only display a reduced level of H3.1K27me1 without major changes to H3.1 deposition ^9^. To examine the impact of H3.1K27me1 deficiency on viral replication, we challenged *fas2* and numerous hypomorphic *H3.1* mutants with CaLCuV. The *fas2* plants showed a more severe yellow mosaic plants than Col-0, whereas the H3.1 single mutants and quadruple mutants showed similar infection ratio and symptom severity to those in Col-0 (Extended Data Fig. 3). Of note, inactivation of *FAS2* or *H3.1* genes did not result in any defect on heterochromatic DNA stability due to the concurrent loss of H3.1K27me1 and H3.1 ^9, 15^. These results imply that concomitant reduction of H3.1K27me1 and H3.1 does not mimic the suppression of viral DNA replication observed in *atxr5 atxr6* where only H3.1K27me1 is reduced.

### Viral resistance of *atxr5 atxr6* is not directly related with TE re-activation and DNA amplification

The *atrx5 atxr6* mutant has three main molecular phenotypes: TE reactivation, heterochromatin amplification and activation of DNA repair pathways^9, 12, 34^. It has been reported that mutations of *MBD9* or *SAC3B* in *atxr5 atxr6* rescue the loss of H3.1K27me1, and suppress heterochromatin amplification and TE reactivation^12, 14^. By contrast, a mutation of *BRCA1,* a gene involved in replication fork stability and DNA repair, enhances both heterochromatin amplification and TE reactivation in *atxr5 atxr6*^12, 14^. We revisited these experiments using leaves #1-#6 of five-week-old mock-treated plants and obtained similar results as the previous report, which used cotyledons in their experiments^12^ (Fig 1c,d). These triple mutant lines provided an opportunity to investigate the relevance of distinct molecular phenotypes of *atxr5 atxr6* in relation to geminiviral pathogenesis in plants.

We inoculated *mbd9*, *sac3b*, *brca1*, *atxr5 atxr6*, *mbd9 atxr5 atxr6*, *sac3b atxr5 atxr6, brca1 atxr5 atxr6* and Col-0 with CaLCuV. The ratio of symptomatic plants and the viral titer in *mbd9*, *sac3b* and *brca1* mutants were comparable to those in Col-0 and higher than those in *atxr5 atxr6*. Intriguingly, *mbd9 atxr5 atxr6*, *sac3b atxr5 atxr6* and *brca1 atxr5 atxr6* mutants all showed a similar ratio of symptomatic plants relative to Col-0, and a significantly higher percentage of symptomatic plants with severe chlorosis compared to *atxr5 atxr6* (Fig. 1e-g, and Extended Data Fig. 4a). Consistently, titers of viral DNA in the three triple mutants were comparable to those in Col-0 and higher than the amount in *atxr5 atxr6* (Fig. 1g). Thus, the susceptibility of *brca1 atxr5 atxr6*, *mbd9 atxr5 atxr6* and *sac3b atxr5 atxr6* mutants was attributed to effect of *BRCA1*, *SAC3B* and *MBD9* on molecular features in *atxr5 atxr6*, but not to their distinct functions in the wild-type condition. Given that three triple mutants had similar viral infection profiles, thus, heterochromatin amplification and TE reactivation were not directly responsible for Geminivirus resistance in *atxr5 atxr6* mutants. In lines with this result, *suvh4/5/6* and *drm1 drm2 cmt3* mutants that have TE reactivation ^34^ showed hyper-susceptibility to viral infection (Extended Data Fig. 1).

### Viral resistance of *atxr5 atxr6* is coupled with the enhanced expression of genes involved in DNA repair

To pinpoint the genetic pathways that attributed to viral resistance of *atxr5 atxr6*, we mined public RNAs-seq data of *mbd9 atxr5 atxr6*, *sac3b atxr5 atxr6*, *atxr5 atxr6* and Col-0 from a cotyledon stage to perform transcriptome-wide association studies (TWAS)^12^. Mutations of *MBD9* and *SAC3B* suppress enhanced expression of 240 protein coding genes in *atxr5 atxr6* (Extended Data Fig. 4c, d). Gene ontology (GO) analysis reveals that significant enriched biological processes belonged to innate immune response, response to salicylic acid (SA) and repair of DSBs (Extended Data Fig. 4e). These three pathways might individually or synergistically contribute to viral resistance of *atxr5 atxr6*.

To investigate how the transcriptome is reprogramed upon virus infection, we performed comprehensive TWAS with high quality reads (Extended Data Fig. 5). When the samples from mock and virus inoculation treatments were considered, approximately 4800 differentially expressed genes (DEGs, fold change ≥2, FDR adjusted *p*-value ≤0.05) were recovered (Extended Data Fig. 6a-c). Among the DEGs, 1136 genes showed enhanced expression upon virus infection (Extended Data Fig. 6d-f). Among the virus infection-activated genes, we selected 365 genes that were expressed at higher levels in *atxr5 atxr6* compared to Col-0, *brca1 atxr5 atxr6*, *mbd9 atxr5 atxr6* and *sac3b atxr5 atxr6*, in both mock-treated and virus-infected samples (Fig. 2a). GO analysis classified the top three enriched biological processes into response to DNA damage (19), DNA repair (17) and DNA recombination (12) (Fig. 2b). Of note, the DNA repair genes were upregulated in Col-0 upon viral infection and then further enhanced in *atxr5 atxr6*; and these genes belong to HRR rather than NHEJ (Fig. 2c). Overall, TWAS revealed a significant association between viral resistance of *atxr5 atxr6* and enhanced expression of DNA repair genes.

**Fig. 2.**
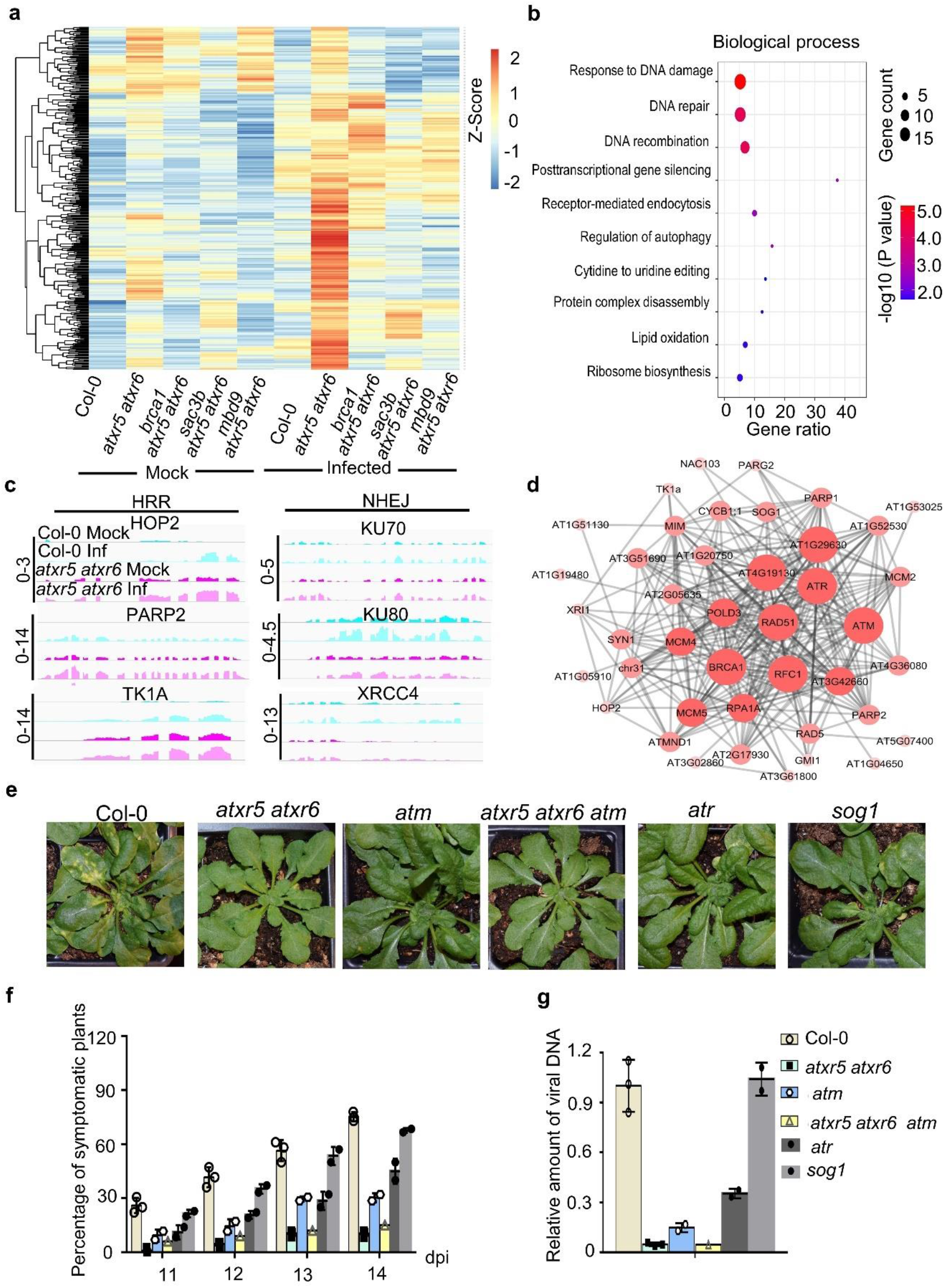
Homology directed repair (HDR) pathway contributes to viral resistance of *atxr5 atxr6*. **a,** Heat map shows the accumulation change of 365 transcripts, selected a total of 4800 DEGs based on the clustering analysis, in mock-treated and virus-infected Col-0, *atxr5 atxr6*, *sac3b atxr5 atxr6* and *mbd9 atxr5 atxr6*. The quantification is conducted by de-seq2. **b,** Bubble plots from Gene Ontology (GO) analysis show the enrichment of 365 genes from (A) in different biological processes. **c,** IGV files show changes of transcript levels of indicated genes in mock-treated and viral-infected Col and *atxr5 atxr6* (Normalized by RPKM). Scales for the distinct loci were shown in left as solid lines. **d,** Protein-protein interaction (PPI) network of DNA repair-related proteins encoded by DEGs upon the virus infection. **e,** Representative phenotypes of CaLCuV-infected Col-0, *atxr5 atxr6*, *atm*, *atxr5 atxr6 atm*, *atr* and *sog1* plants. Photographs were taken at 15 dpi. **f,** Percentages of symptomatic plants induced by CaLCuV infection in indicated backgrounds at 11, 12, 13 and 14 dpi. Each dot in the bar plot represents one replicate, experiments were performed with 36 plants/replicate. **g,** q-PCR shows the amount of viral DNA A in CaLCuV-infected plants in indicated backgrounds at 15 dpi. The relative amount of viral DNA A was first normalized to tubulin DNA control, and then to that of Col-0 where the mean was arbitrarily assigned a value of 1 with standard deviation (SD) from three biological replicates.

### DDR factors are required for efficient amplification of viral genome

In our study, virus infection activated the expression of 53 genes related to DNA damage response (DDR) in *atxr5 atxr6*. Among them, Ataxia-telangiectasia mutated (ATM) and ATR are the kinases that redundantly associate with the majority of DDR factors activated during Geminivirus infection (Fig. 2d). To decipher the relationship between geminiviral replication and DDR activation in plants, we performed virus infection assays with Col-0, *atm*, *atr*, *sog1*, *atxr5 atxr6* and *atxr5 atxr6 atm* plants. Unlike *sog1*, *atm* and *atr* showed reduced ratios of symptomatic plants, milder symptoms and less viral DNA compared to Col-0, but the two single mutants were more susceptible to viral infection than *atxr5 atxr6* (Fig. 2e, f). Remarkably, the ratio of symptomatic plants and amount of viral DNA were comparable between *atxr5 atxr6 atm* and *atxr5 atxr6* (Fig. 2f, g). This epistatic phenotype suggests that *atm* and *atxr5 atxr6* function in the same pathway in relation to viral DNA replication.

### Rep recruits DNA repair proteins to facilitate viral DNA replication

Since ATM is related to HRR, we hypothesized that some HRR factors might directly participate in viral propagation. To test this, we conducted yeast two hybrid (Y2H) screening of 20 selected DDR factors using Rep (Extended Data Fig. 7a), the essential viral replication protein, as bait. Y2H screening recovered RPA1A (replication protein A 1A), RAD51 and PCNA1 as binding partners of Rep (Fig. 3a and Extended Data Fig. 7c). We validated the interaction of Rep with RPA1A and RAD51 through co-immunoprecipitation (Co-IP) experiments using transiently expressed proteins in *N. benthamiana* (Fig. 3b). We hypothesized that if RPA1A and RAD51 were recruited to viral DNA to facilitate viral replication, they should associate with the viral minichromosome. Indeed, chromatin immunoprecipitation-qPCR (ChIP-qPCR) readily detected the enrichment of RPA1A and RAD51 on the viral genome (Fig. 3c).

**Fig. 3.**
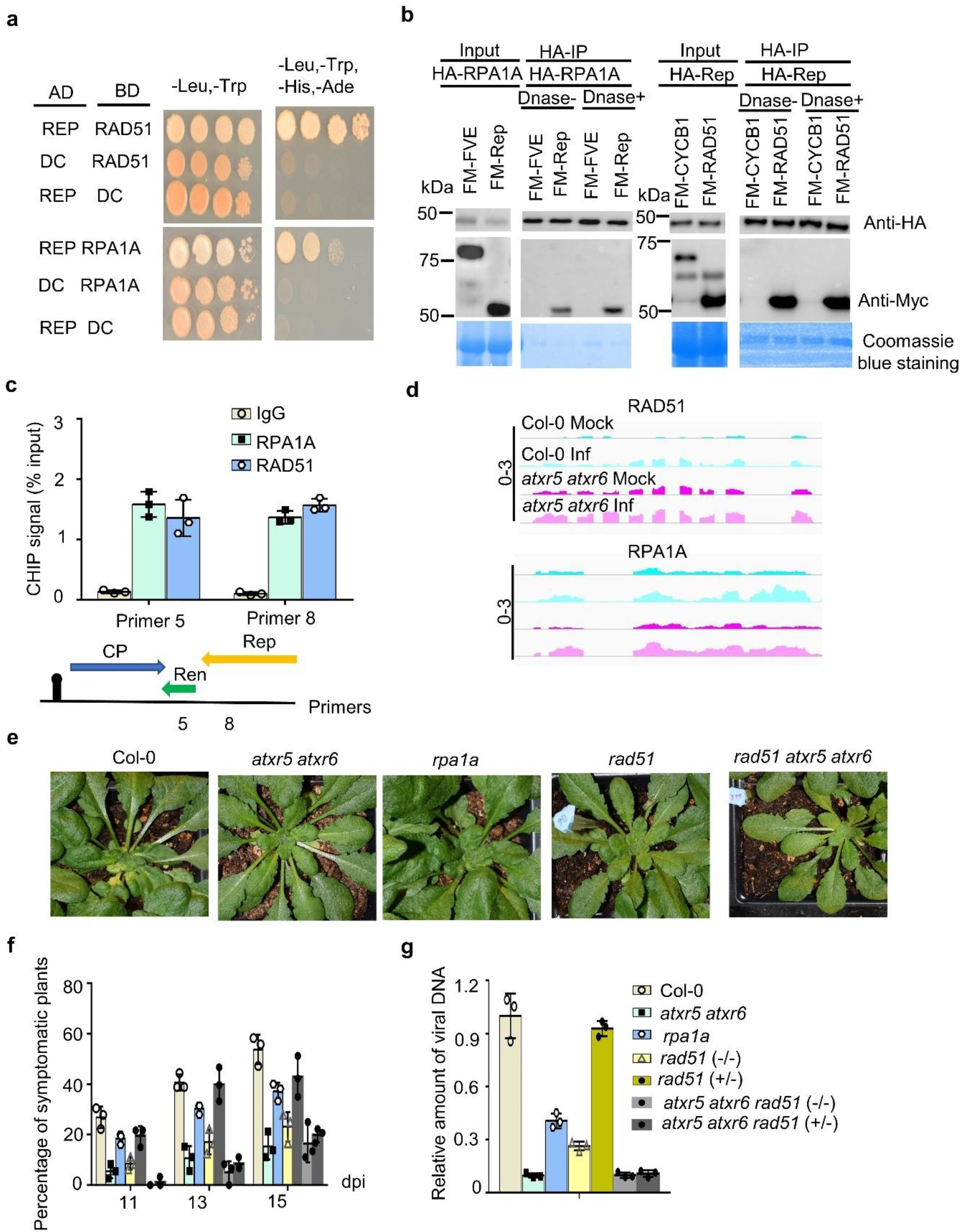
Rep hijacks RAD51 and RPA1A to facilitate viral DNA replication. **a,** Y2H screening pinpointed RAD51 and RPA1 as targets of Rep protein. The negative control is AD/BD vectors. At least 15 independent colonies for each combination were tested and showed similar results. **b,** Co-IP assay validated interactions of Rep with RAD51 and RPA1A in planta. FM-FVE ^72^ and Coomassie blue staining of blots serve as negative and loading controls. **c,** ChIP-qPCR assay shows the binding of RAD51 and RPA1A on viral genome. IgG is a negative control. Error bars represent SD of three replicates. **d,** IGV file shows transcript levels of *RAD51* and *RPA1A* in mock-treated or viral-infected Col-0 and *atxr5 atxr6* (Normalized by RPKM). Scales for the distinct loci were shown in left as solid lines. **e,** Representative phenotypes of CaLCuV-infected Col-0, *atxr5 atxr6*, *rpa1a*, *rad51 (-/-), rad51*(+/-) *atxr5 atxr6 rad51* (*-/-*) and *atxr5 atxr6 rad51* (+/-) plants. Photographs were taken at 16 dpi. **f,** Percentages of symptomatic plants induced by CaLCuV infection in indicated backgrounds at 11, 13 and 15 dpi. Each dot in the bar plot represents one replicate, experiments were performed with 36 plants/ replicate. For *rad51*(-/-), experiments were performed with 15 plants/replicate. **g,** q-PCR assays show the amount of viral DNA A in CaLCuV -infected plants indicated at 16 dpi. Normalization of viral DNA was conducted as Fig. 2g.

We also found that expression of *RAD51* and *RPA1A* was upregulated in *atxr5 atxr6* vs WT and further increased upon virus infection in both backgrounds (Fig. 3d). Importantly, mutations in *RAD51* or *RPA1A* resulted in lower ratio of symptomatic plants and reduced viral DNA accumulation compared to Col-0 (Fig. 3e-g). On the other hand, mutations of *CHR31* or *TK1A* did not affect the viral pathogenesis (Extended Data Fig. 7c,d). These results indicate that RAD51 and RPA1 are essential for Geminivirus replication in *Arabidopsis*. Remarkably, loss of *RAD51* did not have an additive effect on the viral resistance phenotype of *atxr5 atxr6* (Fig. 3g). Given that *atxr5 atxr6* and *atxr5 atxr6 rad51* had the same viral resistance phenotype, we concluded that RAD51 and RPA1A were downstream effectors that accounted for reduced viral replication in *atxr5 atxr6*.

### RAD51 and RPA1A preferentially bind to ribosomal DNA (rDNA) and ncRNA loci in *atxr5 atxr6*

Increased DSBs and RAD51 foci have been observed in over-replication associated centers (RACs), a structure observed during remodeling of heterochromatin, suggesting that RAD51 is involved in DNA repair in *atxr5 atxr6*^11^. On the other hand, RAD51 and RPA1A among other HRR factors are hijacked by Rep to participate in virus replication. These facts raised the possibility that Geminivirus might compete with host RACs in *atxr5 atxr6* for a limiting amount of RAD51, RPA1A and other HDR factors.

To test this, we performed genome wide chromatin immunoprecipitation-sequencing (ChIP-seq) for RAD51 and RPA1A in Col-0 and *atxr5 atxr6* used the specific antibodies against the two proteins (Extended Data Fig. 8a-c). Correlation heat map analysis of ChIP-seq datasets showed high reproducibility among replicates, indicative of the reliability of our ChIP-seq (Extended Data Fig. 8d-f). We initially counted total reads which were uniquely mapped to genome or had multiple mapping locations, and observed increased number of peaks in heterochromatin (Extended Data Fig. 9a, b). We observed that a majority of RPA1A-bound loci coincided with RAD51-enriched regions, despite the fact that much less peaks were identified for the RPA1A ChIP-seq (Fig. 4a). These results suggested that the two proteins might coordinate with each other during HRR. Indeed, their physical interaction could be readily validated in our Co-IP experiments (Fig. 4b). These results were also consistent with earlier reports showing that concomitant absence of RPA1A and RPA1C mimics *rad51* phenotypes (i.e., sterile)^35^.

**Fig. 4.**
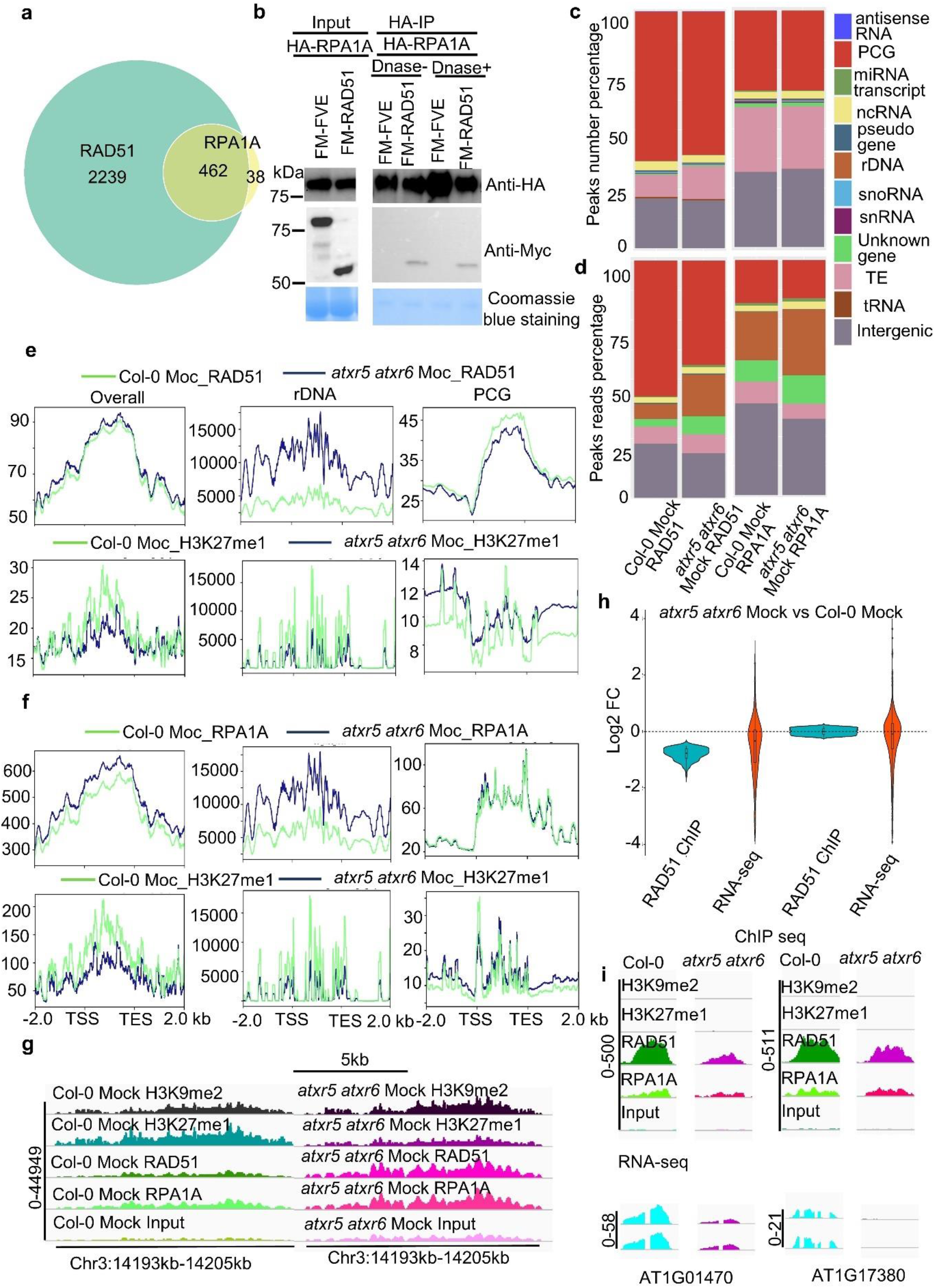
ChIP-seq assays of RAD51 and RPA1A show that unstable genomic DNA reshapes the distribution of HRR events in *atxr5 atxr6* vs Col-0. **a,** Venn graph shows overlap between RAD51 and RPA1A enriched regions in host genomes. **b,** Co-IP assay validated the interaction between RPA1A and RAD51 in planta. FM-FVE ^72^ and Coomassie blue staining of blots serve as negative and loading controls. **c,** Genomic feature classification of RAD51 and RPA1A enriched regions in Col-0 and *atxr5 atxr6*. The y axis represents the peak number percentages of various categories. **d,** Peak density distributions of RAD51 and RPA1A ChIP-seq in Col-0 and *atxr5 atxr6*. The y axis represents the reads percentage of various categories. **e-f,** Distribution of normalized ChIP-signal (RPKM) of RAD51 and RPA1A **f** and H3K27me1 from Col-0 and *atxr5 atxr6* over different categories. H3K27me1 related data were mined from published data (GSE111814). **g,** IGV files of normalized ChIP signals of H3K9me2, H3K27me1 (RPKM), RAD51, and RPA1A on a rDNA locus. H3K9me2 and H3K27me1 related data were mined from published data (GSE111814). Scales for the distinct loci were shown in left as solid lines. **h,** Violin plot shows the expression changes of transcripts from the loci with reduced and unchanged RAD51 ChIP signal in *atxr5 atxr6* vs Col-0. **i,** IGV files of normalized ChIP signals of H3K27me1, RAD51, and RPA1A (RPKM) and normalized transcript levels (RPKM) on selected loci from RNA-seq. Scales for the distinct loci were shown in left as solid lines.

We found that both RAD51 and RPA1A bound to broad heterochromatic regions including rDNA, small nucleolar RNAs (snoRNAs), TEs, and ncRNAs in mock treated Col-0 (Figure 4C). RAD51 and RPA1A were also distributed over euchromatic regions that generate protein coding transcripts (Fig. 4c). Interestingly, this pattern is reminiscent of ChIP-seq patterns of RAD51 and RPA1A in *Mus musculus*^36^, suggestive of their conservative functions in eukaryotes (Extended Data Fig. 9c). We next compared peak numbers in heterochromatic elements in *atxr5 atxr6* and Col-0. The overall patterns of RAD51 and RPA1A-bound peaks over rDNA, ncRNA, and TE among the others seems not to be affected by the loss of *ATXR5/6* relative to Col-0 (Fig. 4c). One possible reason is that peaks numbers over heterochromatic regions represent a relatively small fraction of the total called peaks. We further assessed the peak density profiling of RAD51- and RPA1A-occupied regions by calculating the fractions of total reads for the peaks from different categories. Interestingly, we observed a substantial increase in RAD51 and RPA1A occupancy at the loci corresponding to novel transcripts, rDNA (Fig. 4d-f) and ncRNAs (Extended Data Fig. 10a, b) in *atxr5 atxr6* vs Col-0. In other words, RAD51 and RPA1A displayed a robust increase in read coverage over rDNA among other classes in *atxr5 atxr6* compared to Col-0 (Fig. 4d-f).

Emerging evidence shows the association between *RPA, RAD51, BRCA1* and *RAD51*-associated protein 1 (*RAD51AP1*) with transcription processes. It has been also shown that reduced H3K27me1 on the *45S rDNA* loci induces the expression of *45S rRNA* variants, which is accompanied by higher copy number of *45S rDNA* in 8C nuclei of *atxr5 atxr6* ^37^. In our hands, we found that RAD51 and RPA1A were significantly enriched on two sites of rDNA and the enrichment was well correlated with increased copy number of rDNA (Fig. 4e-g, Extended Data Fig. 10c) in *atxr5 atxr6* compared to Col-0. Furthermore, the increased rDNA content in *atxr5 atx6* vs Col-0 is concomitant with reduced level of H3K27me1 (Fig. 4e-g, Extended Data Fig. 10c). These results suggested that H3K27me1 might acts as a repressive marker to regulate rDNA stability and recruitment of HRR factors onto rDNA. By contrast, reduced H3K27me1 over heterochromatic regions observed in *atxr5 atxr6* was restored in *mbd9 atxr5 atxr6* and *sac3b atxr5 atxr6* mutants ^14^, and also at rDNA loci (Extended Data Fig. 11a). Collectively, these results strongly support a model where unstable DNA in *atxr5 atxr6*, especially rDNA loci, requires RAD51 and RPA1A, among other HRR factors, to maintain genome integrity.

### Coordination between reduced HRR occupy and impaired transcription at defense-related loci in *atxr5 atxr6*

Besides rDNA loci, RAD51 and RPA1A occupied numerous protein coding gene (PCG) loci (Fig. 4c-f). A gene ontology (GO) analysis showed that a large number of RAD51- and RPA1A-enriched genes belonged to several genetic pathways, such as response to cold, salt stress, oxidative stress and jasmonic acid (JA)-mediated signaling (Extended Data Fig. 11b, c). In addition, RAD51 also bound genes related to defense response, response to SA, cell communication, immune system, fatty acid and other important biological processes (Extended Data Fig. 11b). Interestingly, the PCGs enriched in RAD51 or RPA1A showed very low levels of H3K27me1 in Col-0 and *atxr5 atxr6*, supporting an active chromatin status at these loci (Fig. 4e, f). Further analysis showed that the signals of RAD51 and RPA1A tended to be evenly distributed over gene bodies rather than enriched at promoters in Col-0 and *atxr5 atxr6* (Fig. 4e, f). In contrast to *rDNA* loci, RAD51 signal tended to be reduced over the bodies of PCGs in *atxr5 atxr6* vs. Col-0 (Fig. 4e). We then selected 366 PCGs that showed lower RAD51 ChIP signal in *atxr5 atxr6* vs Col-0 (analyzed by de-seq2, log2FC<-0.5, *p*-value<0.05) and assessed their transcript levels. Among the selected genes, the transcripts of 207 PCGs were detectable. Interestingly, 71.0% of the PCGs not only showed decreased RAD51 ChIP signal but also showed reduced transcript accumulation in *atxr5 atxr6* relative to Col-0 (Fig. 4h, i). Thus, our data indicate that reduced occupancy of RAD51 over otherwise actively transcribed regions coincided with their decreased transcript accumulation in *atxr5 atxr6*. The correlation between DNA repair and transcription observed here is reminiscent of a recent discovery of transcription-associated homologous recombination repair (TA-HRR)^38^. As one detrimental byproduct of transcription, unprocessed R-loops often cause the formation of DSBs, which in turn inhibit local ongoing transcription^39^. Supporting the importance of HRR factors in promoting transcription, RAD51 associated protein 1 (RAD51AP1) induces the formation of R-loop and favors RAD51-mediated D-loop formation to restore active transcription over these regions^40^. The fact that normally active chromatin regions are depleted of HRR factors to the point of interfering with transcription suggests that the availability of HRR factors is limiting in *atxr5 atxr6* mutants, despite increased expression of these factors in this mutant background.

### Reduced RAD51 and RPA1A binding to viral DNA in *atxr5 atxr6*

Based on our ChIP-seq results, we hypothesized a competition between host and viral genomes for HRR factors. In *atxr5 atxr6* mutants, HRR factors would be sequestered at unstable DNA regions, preventing them from being used by the viral Rep for pathogen replication. To test this, we compared RAD51 and RPA1A ChIP-seq of mock and virus infected plants. Our initial ChIP-qPCR detected a significant enrichment of RAD51 and RPA1A in viral genome in Col-0 but barely any signal in the *atxr5 atxr6* background when representative samples were randomly collected at an earlier stage (Extended Data Fig. 12a). This result clearly resulted from that the fact that *atxr5 atxr6* had significantly reduced infection rate and viral load than Col-0. To account for differences in virus titer, we arbitrarily increased the number of *atxr5 atxr6* symptomatic plants to artificially mimic the infection ratio of Col-0 for ChIP-seq assays to test distribution of RAD51 and RPA1A among virus and host.

For Col-0, virus infection resulted in reduced signal of RAD51 and RPA1A over host chromatin in virus infected plants vs mock. The decreased signals of RAD51 and RPA1A mainly originated from the PCG loci rather than rDNA regions, likely due to relative lower DNA damage in Col-0 plants vs *atxr5 atxr6* under normal conditions^11^ (Fig. 5a-c, Extended Data Fig. 12b-d). Notably, the genes with reduced RAD51 occupancy upon viral infection were related to well-known defense pathways involving JA, fatty acid and s-adenosylmethionine (SAM) (Extended Data Fig. 13a). Moreover, the steady-state transcript levels from those loci were also reduced in infected Col-0 plants when compared to the mock, prompting the question of whether the Geminivirus genome suppresses the host immune system through the sequestration of RAD51 (Extended Data Fig. 13b).

**Fig. 5.**
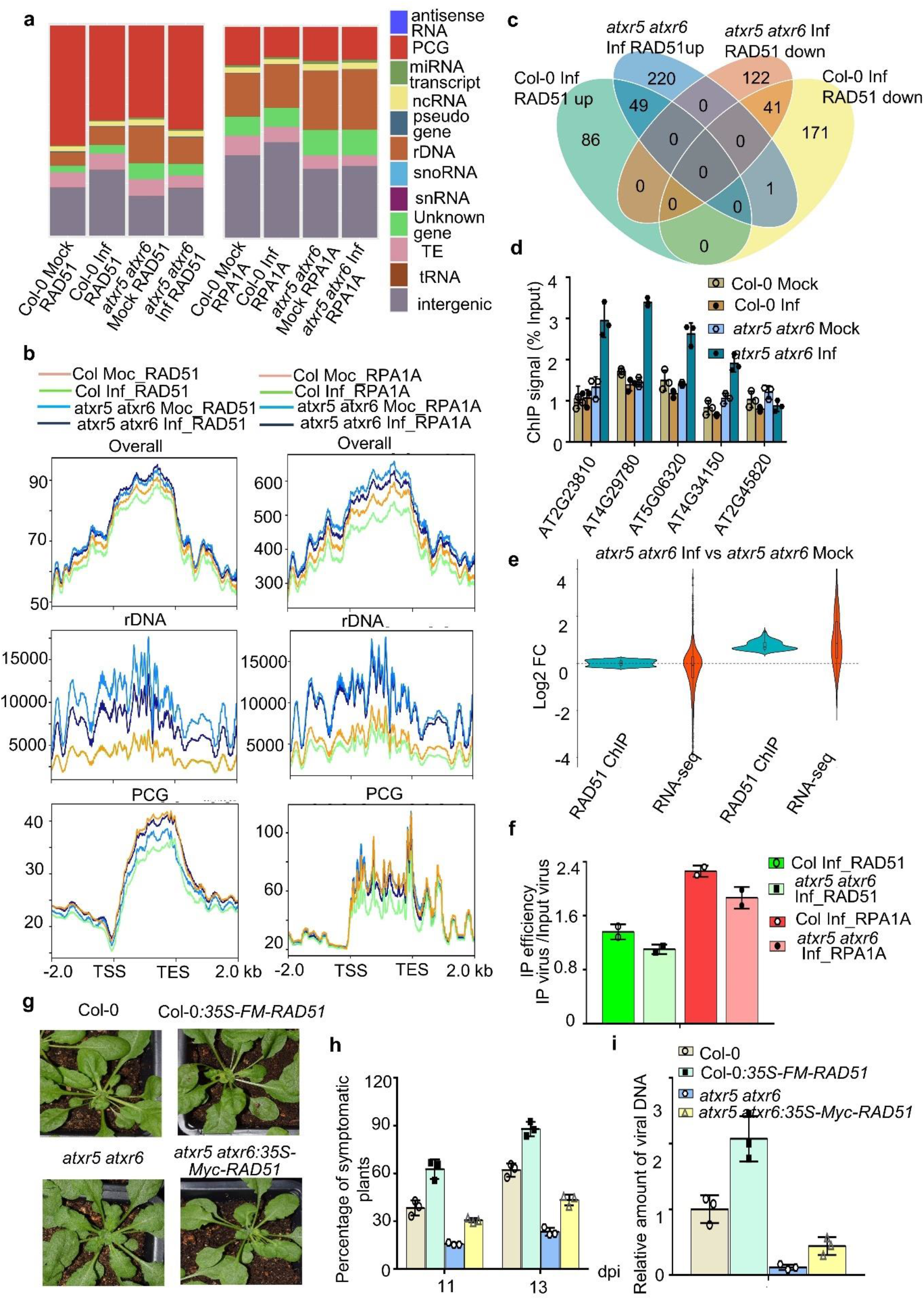
Recruitment of HRR factors onto unstable genomic DNA and defense related genes prevents the loading of HRR factors on viral genome. **a,** Peak density distributions of RAD51 and RPA1A ChIP-seq in various genomic regions from mock-treated and virus-infected Col-0 and *atxr5 atxr6*. **b,** Distribution of normalized ChIP signal of RAD51 and RPA1A (RPKM) in different genome categories mock-treated and virus-infected Col-0 and *atxr5 atxr6*. **c,** Venn graph shows change of RAD51 signal from enriched peaks over host genome in Col-0 and *atxr5 atxr6* upon virus infection. **d,** ChIP-qPCR shows that significantly increased RAD51 binding of host genome in *atxr5 atxr6* on selected loci. *AT2G45820* serves as a negative control. **e,** Violin plot shows expression changes of transcripts from the loci with unchanged and increased RAD51 ChIP signal in viral-infected vs mock-treated *atxr5 atxr6*. **f,** Loading efficiency of RAD51 and RPA1A on viral genome in infected Col-0 and *atxr5 atxr6*. Loading efficiency was calculated from the percentage of reads mapped to virus genome in the ChIP-seq of RAD51 and RPA1A relative to the ones from corresponding inputs. **g,** Representative pictures of the virus-infected plants in indicated lines. Photographs were taken at 14 dpi. **h,** Percentages of symptomatic plants of viral-infected plants in indicated backgrounds at 11 and 13 dpi. Each dot in the bar plot represents one replicate, experiments were performed with 36 plants/replicate. **i,** Constitutive expression of RAD51 promotes the viral DNA replication in Col-0 and *atxr5 atxr6*. qPCR assays show increase of virus titers in RAD51 overexpression lines vs their reference backgrounds. Normalization of viral DNA was conducted as Fig. 2g.

In contrast to Col-0, RAD51 ChIP signal displayed a moderate increase whereas RPA1A showed minor decrease in host genome of *atxr5 atxr6* upon the virus infection vs mock treatment (Fig. 5b). Detailed plotting of ChIP signals showed the binding of RAD51 decreased over rDNA and ncRNAs loci but increased over the PCGs in *atxr5 atxr6* upon virus infection relative to the mock (Fig. 5b and Extended Data Fig. 12b). Notably, we found viral infection specifically enhanced the RAD51 signal over 270 PCGs in *atxr5 atxr6* whereas only 18.1% of them showed increased RAD51 signal in Col-0 (Fig. 5c, d). Among 270 PCGs, transcripts of around 177 PCGs were detectable, 78.5% of 177 PCGs also displayed accumulated transcripts in the infected *atxr5 atxr6* plants (Fig. 5e). These genes are mainly related to defense and immune responses, SA, and JA, indicating that the virus infection enhances the RAD51 signal and associated immune response exclusively in *atxr5 atxr6* (Extended Data Fig. 13d). Importantly, ChIP signals of RAD51 and RPA1A were significantly enriched over both heterochromatin (rDNA) and PCGs in the infected *atxr5 atxr6* mutants compared to those of infected Col-0 plants (Fig. 5b). These results suggested that the occupancy of RAD51 and RPA1A in the heterochromatic regions and PCGs both contributed to the viral resistance of *atxr5 atxr6* vs Col-0.

The prediction of our model is that the loading of RAD51 and RPA1A on viral genome is reduced in *atxr5 atxr6* vs Col-0. Indeed, the loading efficiency of RAD51 and RPA1A on the viral genome, reflected by the ratio of reads mapped to virus from IP samples divided by the numbers from input samples, was reduced in *atxr5 atxr6* vs Col-0 (Fig. 5f). This result clearly indicated that even RAD51 and RPA1A accumulated in genome of *atxr5 atxr6* but could not be efficiently loaded to the viral minichromosome for replication upon viral infection. Collectively, loss of ATXR5/6 elevated RAD51 and RPA1A signals on host genome and prevented their loading onto the viral genome, leading to the resistance phenotype of *atxr5 atxr6*.

To further test our model, we generated Col-0:*35S-FM-RAD51* and *atxr5 atxr6:35S-MYC-RAD51* and inoculated the stable transgenic lines with the virus (Extended Data Fig. 8a). Indeed, overexpression of RAD51 in both Col-0 and *atxr5 atxr6* clearly increased the ratio of symptomatic plants and viral DNA amount compared to their own controls (Fig. 5g-i). Altogether, we concluded that re-replicated DNA and the loci of defense-related PCGs in *atxr5 atxr6* take up HRR components, leading to shortage of the components essential for viral replication in *atxr5 atxr6* vs Col-0.

### BRCA1, HOP2 and CYCB1 are *bona fide* partners of RAD51

We next aimed to identify the factors that potentially recruited RAD51 and RPA1A to the host genome. Through Y2H assays, we recovered BRCA1 and B1 type cyclin dependent protein kinase CYCB1;1 as partners of RAD51 from five candidate genes (Extended Data Fig. 14a). Neither protein appeared to directly interact with RPA1A in the assays (Figure 6A). RAD51 could be readily detected in the co-immunoprecipites of CYCB1;1 and BRCA1 but not in that of the corresponding negative controls (Fig. 6b). In addition, Homologous pairing protein 2 (HOP2) interacted with RAD51 in Co-IP, but not in Y2H assay (Fig. 6b, Extended Data Fig. 14a). Of note, all three proteins are intrinsically disordered or contain disordered segments; and these features might contribute to their interaction with RAD51 (Extended Data Fig. 14b).

**Fig. 6.**
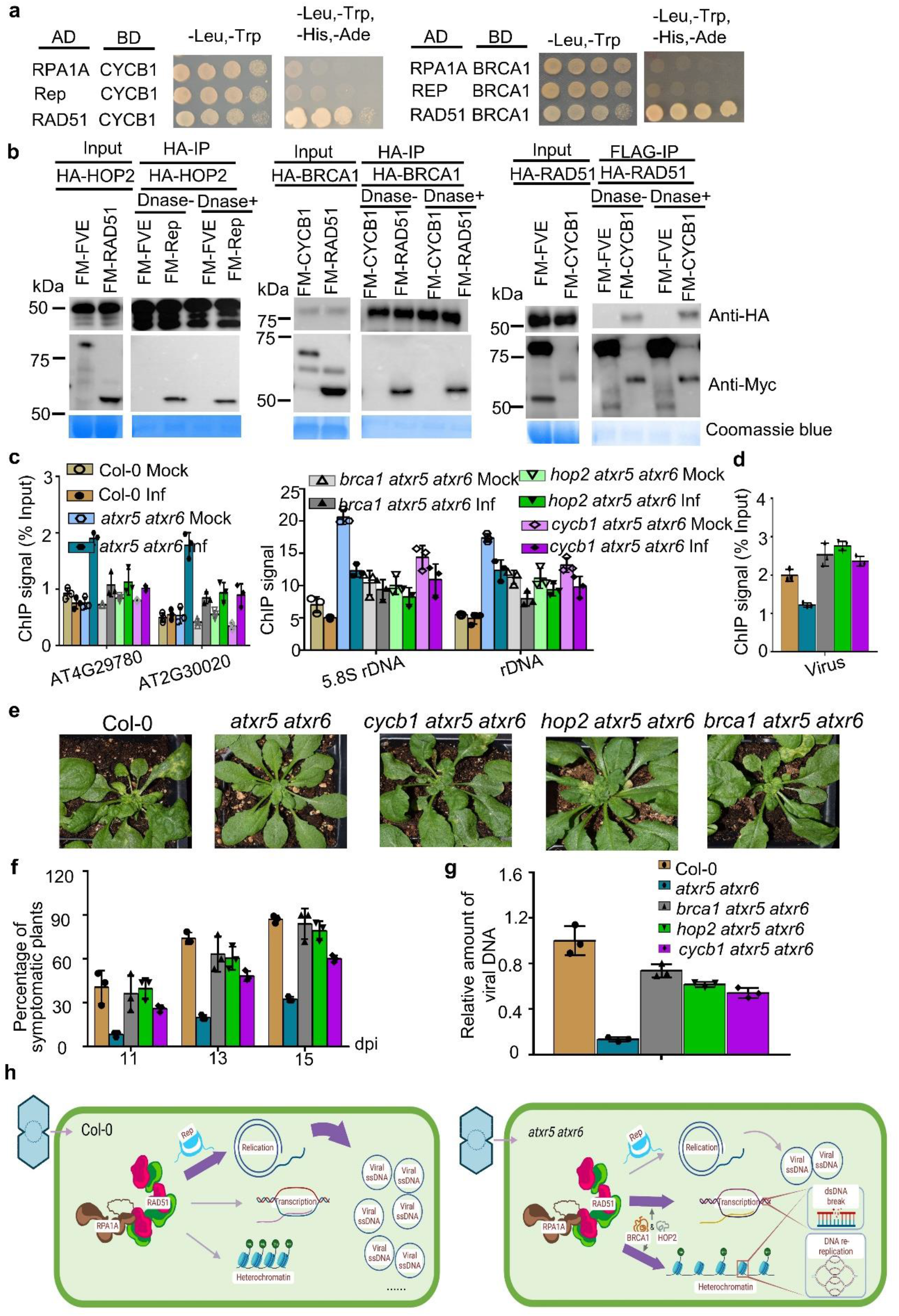
BRCA1, HOP2 and CYCB1 regulate the antiviral defense in an RAD51-dependent manner. **a,** Y2H assays validated the interactions of RAD51 with CYCB1 and BRCA1. **b,** Co-IP assay validated the interactions of RAD51 with CYCB1, BRCA1, and HOP2 in planta. FM-FVE ^72^ and Coomassie blue staining of blots serve as negative and loading controls. **c,** ChIP-qPCR assay shows that BRCA1, HOP2 and CYCB1 are required for recruitment of RAD51 over rDNA and PCGs in *atxr5 atxr6*. The ChIP signal of *rDNA* was normalized to that of AT1G40104. **d,** ChIP-qPCR assay shows that BRCA1, HOP2 and CYCB1 are required for reduced binding of RAD51 over viral genome in *atxr5 atxr6*. **e,** Representative pictures of the virus-infected plants in indicated backgrounds. Photographs were taken on at 15 dpi. **f,** Percentages of symptomatic plants of viral-infected plants in indicated backgrounds. Each dot in the bar plot represents one replicate, experiments were performed with 36 plants/replicate. **g,** q-PCR assays show the amount of viral DNA A in CaLCuV -infected plants indicated at 16 dpi. Normalization of viral DNA was conducted as Fig. 2g**. h,** A proposed model for Geminivirus competition with host genome for HRR to facilitate viral replication. In the healthy WT with stable genome, the limited amount of DNA repairing proteins is distributed in the heterochromatic regions and deposited onto the defense-related genes. Thus, the viral-encoded Rep protein could hijack RAD51 and RPA1A for efficient viral replication. However, in *atxr5 atxr6*, unstable host DNA and defense-related genes retain a large amount of DNA repairing machinery to prevent its routing to viral genome, leading to low efficient replication of the virus.

### Roles of BRCA1, HOP2 and CYCB1 in loading RAD51

Interactions of BRCA1, HOP2 and CYCB1;1 with RAD51 detected here and also in mammalian cells and plants^41–43^ suggested the potential roles of these proteins in recruitment of RAD51 onto unstable DNA. Robust increase of viral DNA was detected in *brca1 atxr5 atxr6* compared to that in *atxr5 atxr6,* and no obvious difference in the viral titer was detected between *brca1* and Col-0 (Fig. 1e-g). These contrasting results suggested that RAD51-centered DNA repair components were not able to reach the unstable loci in *atxr5 atxr6*; rather, the components were re-routed to viral minichromosome for DNA replication in *brca1 atxr5 atxr6* compared to *atxr5 atxr6* (Fig. 6c, d). Differently, DNA repair components were largely recruited to viral genome in the single mutant *brca1* and Col-0 where host DNA is intact in a normal physiological condition. Thus, BRCA1 might regulate viral DNA replication in a manner dependent on RAD51 and other HRR factors in the *atxr5 atxr6* background.

To test whether deletions of *HOP2* and *CYCB1;1* would also interrupt recruitments of HRR factors between host DNA and viral genome in *atxr5 atxr6*, we generated *cycb1 atxr5 atxr6* and *hop2 atxr5 atxr6* mutants through genetic crossing and tested their susceptibility to viral infection. When challenging the higher order mutants with CaLCuV, *hop2 atxr5 atxr6* and *cycb1 atxr5 atxr6* mutants, like *brca1 atxr5 atxr6,* had higher percentages of symptomatic plants and more severe chlorosis phenotype than those in *atxr5 atxr6* (Fig. 6e-g). Moreover, *hop2 atxr5 atxr6* and *cycb1 atxr5 atxr6* mutants showed higher viral DNA content relative to *atxr5 atxr6*, but lesser than that of *brca1 atxr5 atxr6* (Fig. 6e-g). These results indicated that BRCA1, HOP2 and CYCB1;1 were essential for the virus resistance phenotype in *atxr5 atxr6*, with suggestion that the three proteins might be responsible for recruitment of RAD51 to the unstable loci and PCGs in the host genome. Of note, differences between the triple mutants were also observed (Fig. 6e-g), suggesting that their contributions to the loading of RAD51 or other HRR factors onto damaged DNA might be different. Alternatively, additional paths for recruitment to RAD51 and other HRR factors might compensate for the loss of the three components in different degrees.

To further test our model, we selected several representative loci on heterochromatic regions and the PCGs to perform ChIP-qPCR. Indeed, RAD51 signals over heterochromatic regions including 5.8S rDNA were all lower in the triple mutants than that in *atxr5 atxr6* under mock treatment and infection conditions. This result indicated the deficient recruitment of RAD51 onto host heterochromatic regions in the triple mutants vs *atxr5 atxr6* (Fig. 6c). Quantitative analysis showed that RAD51 occupancy on the above loci in mock treated *cycb1 atxr5 atxr6* was higher than that in *hop2 atxr5 atxr6* and *brca1 atxr5 atxr6*, suggesting that CYCB1;1 might contribute less to recruitment of RAD51 vs BRCA1 and HOP2 proteins. Moreover, deletions of *CYCB1;1*, *HOP2* and *BRCA1* in *atxr5 atxr6* also prevented the loading of RAD51 on PCGs upon virus infection (Fig. 6c). On the other hand, more efficient loading of RAD51 on viral genome was observed in *hop2 atxr5 atxr6*, *brca1 atxr5 atxr6* and *cycb1 atxr5 atxr6* compared with that of *atxr5 atxr6* (Fig. 6d). In contrast, in WT background where there was barely HRR and the amount of RAD51 was sufficient for viral replication, deletions of *CYCB1;1*, *HOP2* and *BRCA1* did not affect the viral pathogenesis and accumulation (Extended Data Fig. 14). Thus, we concluded that BRCA1, HOP2 and CYCB1;1 indirectly regulate viral DNA replication through controlling the availability of RAD51 to the viral genome in the *atxr5 atxr6* background (Fig. 6h).

## Discussion

Here we reported a novel regulatory role of histone MTases on DNA virus amplification. Different from other histone MTases that deposit repressive marks on viral genomes and typically repress viral infections, ATXR5/6 promote viral replication as a trade-off to maintaining host genome integrity. This unique mode of action in viral infection is highlighted by the fact that loss-of-function mutants of *atxr5 atxr6* become resistant to the virus compared to wild-type Col-0. The underlying mechanism for the viral resistance is that mutations of *ATXR5* and *ATXR6* causes unstable heterochromatic regions exemplified by rDNA and ncRNAs loci, which activate the HRR pathway. These unstable heterochromatic regions together with defense related PCGs in euchromatin regions in turn sequester HRR factors and preclude them from being hijacked by the virus-encoded Rep protein, leading to suppression of the viral replication (Fig. 6h). Several pieces of evidence support our model: 1) DNA repair factors such as ATM and ATR are required for efficient viral DNA replication (Fig. 2e-2g); 2) the viral protein Rep hijacks RAD51 and RPA1A among other DNA repair components to facilitate viral replication (Fig. 3c-g); 3) although *MBD9*, *SAC3B*, and *BRCA1* play opposite roles in regulating heterochromatin amplification and TE reactivation, loss of any of these genes interrupts activation of HRR and restore viral replication in *atxr5 atxr6* to similar levels observed in Col-0. These observations suggest that heterochromatin amplification and TE reactivation in *atxr5 atxr6* do not function as a prophylactic system against CaLCuV infection (Fig. 1c-g); 4) reduced H3K27me1 causes extra copies of heterochromatic DNA, including rDNA and ncRNA loci, which induces the recruitment of HRR factors such as RAD51 and RPA1A at heterochromatin loci in the host (Fig. 4d-g); 5) deficient loading of RAD51 and RPA1A on viral genome is detected in *atxr5 atxr6* relative to Col-0, despite higher accumulation of the proteins in the mutant and stronger ChIP signal over the *atxr5 atxr6* genome relative to Col-0 (Fig. 5b-5f); and 6) BRCA1, HOP2, and CYCB1;1 promote the recruitment of RAD51 onto the host genome, whereas depletions of *HOP1*, *BRCA1* and *CYCB1* in *atxr5 atxr6* caused inefficient loading of RAD51 onto the unstable host genome and PCGs related to defense, and re-routing of RAD51 onto viral genome in *atxr5 atxr6*, leading to hyper-susceptibility of viral infection in the triple mutants vs *atxr5 atxr6* (Fig. 6c-g). Thus, we conclude that RAD51 and RPA1A, among other HRR factors, are the cornerstones in the battle of host and virus. If the virus fails to recruit RAD51 and RPA1A, it will fail to parasitize the host, as seen in *atxr5 atxr6*.

DSBs activate HRR to faithfully repair DNA during replication to maintain genome stability ^44^. As parts of heterochromatic elements, rDNA is considered a hot spot for recombination events due to their repetitive elements. During HRR, the RPA complex coats resected ends to prevent their degradation. RAD51 then replaces RPA and mediates the D-loop formation and strand invasion ^44^. Robust loading of RAD51 and RPA1A onto rDNA and ncRNAs loci accompanies lower H3.1K27me1 levels in *atxr5 atxr6* (Fig. 4e, f). Similarly, reduced H3.1K27me1 correlates with higher rDNA and ncRNA copy number (Fig. 4e, Extended Data Fig. 10c). These results suggest that RAD51 and RPA1A participate in maintaining genome stability at heterochromatic regions in the absence of H3.1K27me1. In our study, BRCA1 and HOP2 promote the recruitment of RAD51 over rDNA regions to facilitate DNA repair in *atxr5 atxr6*. A similar mechanism has been reported in mammalian cells, where BRCA1 and HOP2 complexes stabilize the RAD51-ssDNA filament and promote RAD51-mediated homologous DNA pairing process in vitro^41, 45^. In *Arabidopsis*, CYCB1-CDKB1 complex phosphorylates RAD51 in vitro^42^. Phosphorylation of RAD51 is required for its efficient DNA binding^46^. In our study, CYCB1 could also impact the recruitment of RAD51 over rDNA, but less efficiently compared to BRCA1 or HOP2, implying the presence of a yet unidentified kinase(s) that target RAD51. Remarkably, loss-of-function mutations of *BRCA1, HOP2*, *and* CYCB1 in *atxr5 atxr6* restored viral resistance to different extents (Fig. 6c-g), suggesting robust loading of RAD51 over heterochromatic region in *atxr5 atxr6* mutants is required for its viral resistance.

RAD51 is also distributed along the gene body of PCGs with low H3K27me1 under physiological conditions (Figure 4E). Furthermore, decreased RAD51 ChIP signals over PCGs is concordant with reduced transcripts levels in *atxr5 atxr6* (Fig. 4h). Notably, the RAD51-enriched PCGs are dedicated to the defense response, response to hypoxia, SA, JA and cold and salt stresses. These observations suggest that RAD51 might coordinate TA-HRR to maintain genome integrity, while coping with physiological stresses (Extended Data Fig. 11b). Exogenous SA treatment induces DNA breaks and promotes RAD51 binding to the promoters of pathogenesis related genes (PR). Conversely, depletion of BRCA2A^27^ and ATR^28^, two partners of RAD51, suppresses the expression of SA-induced defense related genes and efficient immune response. These results further imply that RAD51 is required for efficient transcription regulation and defense response. Importantly, the fact that PCGs display reduced occupancy of RAD51 while producing lower expression of defense-related transcripts in infected Col-0 plants implies a dual role of Rep during infection (Extended Data Fig. 13a, b): hijacking the HRR factors to facilitate the viral genome amplification whereas attenuating the plant defense system in WT plants. By contrast, RAD51 occupancy over defense related genes and transcription of the corresponding loci were enhanced in *atxr5 atxr6* upon virus infection (Fig. 5c-e, Extended Data Fig. 13d), implying that defense genes in *atxr5 atxr6* compete with the virus for a limited amount of RAD51, restricting virus replication and enhancing the immune response, thus resulting in viral resistance.

During the parasitic lifestyle, DNA viruses activate the host DNA repair mechanism and hijack the replication machinery to the viral genome. Here, *rad51* and *rpa1a* mutants displayed resistance to Geminivirus infection, indicating that the proteins are critical for viral replication. Consistently, RAD51 interacts with the Rep encoded by mungbean yellow mosaic India virus (MYMIV) and promotes geminiviral DNA replication in the heterologous system *Saccharomyces cerevisiae*^47^. Geminiviruses replicate their genomes through rolling circle replication and recombination dependent replication^2^. RAD51 might function as a recombinase to directly facilitate the rolling circle replication of the virus. RPA1A might also contribute to the rolling circle replication of geminiviruses, reminiscent of RPA-mediated enhancement of the replication of simian virus 40^48, 49^. As key components for homologous recombination, RAD51 and RPA1A might also directly promote the recombination dependent replication of Geminiviruses. Supporting of this model is that RAD51D, a paralog of RAD51, has been reported to promote the recombination dependent replication of Geminivirus^50^. In our studies, although viral infection enhances the expression of RAD51 and RPA1A in Arabidopsis, the ChIP signals of RAD51 and RPA1A over the host genome were reduced in infected vs mock treated Col-0 plants (Figs. 3d and 5b). Furthermore, Rep interacts with RAD51 and RPA1A in planta (Fig. 3b). All these observations clearly indicate that Rep hijacks RAD51 and RPA1A to facilitate viral amplification during the infection process.

In summary, we propose that robust retention of RAD51 and RPA1A onto unstable host DNA, along with increased RAD51 accumulation over defense-related PCGs, prevent the efficient loading of the replication-essential factors onto the viral genome, leading to a resistance phenotype in *atxr5 atxr6*. This study implies that the virus might adapt to the healthy host to hijack host DNA repairing components for the viral replication. On the other hand, the host could retain DNA repairing proteins onto unstable genome via compromising H3.1K27me1 methyltransferase functions but gain its resistance to viral infection in plants. One implication of this study is that one might apply a certain level of genome toxicity stress on crops or insert certain pieces of RAD51-favored unstable DNA elements into crop genomes^51^. Such actions might prime local DNA repair and trap HRR factors onto host genomes to a certain degree, while granting viral resistance, ensuring agricultural production. Similarly, disabling H4K20me1 transferases in human, the functional homologs of ATXR5/6, can also activate and TONSOKU-LIKE and BRCA1 machinery^52, 53^. Host retention of the DNA repairing machinery to correct genome instability would be expected to increase host immunity and impair the DNA virus amplification, thus providing an opportunity to defend against the viruses. In addition, breaks on rDNA lead to the loss of rDNA repeats and exacerbate the genomic instability during aging^54, 55^. Overexpression of RAD51 restored the accumulated DSBs in aged cells and extended the replicative life span of yeast^55, 56^, implying the potential application of H4K20 and RAD51 on curing rDNA associated human diseases such as amyotrophic lateral sclerosis, and Huntington’s disease^57^.

## Acknowledgements

We thank D. Shippen for *atm*, *atr*, *sog1* seeds. The work was supported by grants from the NIH (GM127742) to X.Z; NIH (R35GM130272) to S.E.J.; Z.W., C.Z., and K.X. were partially supported by China Scholar Council fellowships; S.E.J. is an investigator of the Howard Hughes Medical Institute.

## Author contributions

X.Z. conceived the project. Z.W., CC-G., and X.Z., designed the experiments. Z.W. performed the experiments, conducted the bioinformatic analysis for RNA-seq. and analyzed the majority of data. CC-G. performed the early viral pathogenesis experiments and offered guidance and suggestion through the study. C.Z. performed the initial viral pathogenesis screening of epigenetic related mutants. Z. L. genotyped mutants and validated the viral resistance phenotype. K.X. helped to generate high order mutants. C.T. conducted the bioinformatic analysis for ChIP-seq with the guidance of C.L J. Z. helped RNA-seq library construction and drew the model. Z.S.W., S.E.J., S.D.M., and Y.J., obtained a serial of epigenetic mutants and provided intellectual advice. Z.W. wrote the initial draft of manuscript. X.Z thoroughly edited the paper.

## Methods

### Plant materials and growth condition

*A. thaliana* plants were grown under cool-white, fluorescent light (120 μmol m^-2^ s^-1^) in a short day condition (8 h light/ 16 dark) with 22℃ and 50% humidity. Col-0 was used as a wild type and *atxr5 atxr6, atxr5, atxr6, mbd9, mbd9 atxr5 atxr6, sac3b, sac3b atxr5 atxr6, brca1, brca1 atxr5 atxr6, atm, atr, sog1, atm atxr5 atxr6, fas2, htr1, htr2, htr3, htr9, htr1 htr2 htr3 htr9, suvh4 suvh5 suvh6* were described previously^9, 11, 12, 15, 34^. Seeds of *drm1 drm2 cmt3* (CS16384), and *clf-28* (SALK_139371) were obtained from Arabidopsis biological center (ABRC). T-DNA insertion mutants of *CYCB1 (SALK_200647), HOP2 (SALK_136002), CHR31 (SALK_204501), TK1A (SALK_097767), RAD51 (SAIL_873_C08), RPA1A (SALK_017580)* were purchased from ABRC and genotyped by PCR. High order mutants *cycb1 atxr5 atxr6*, *hop2 atxr5 atxr6* and *rad51* (+/-) *atxr5 atxr6* were generated by genetic crossing. The triple mutant of *rad51* (-/-) *atxr5 atxr6* was generated from selfed *rad51* (+/-) *atxr5 atxr6* and genotyped by PCR before the virus infection assays due to the sterility of *rad51 (-/-)*. For *cycb1 atxr5 atxr6*, *hop2 atxr5 atxr6*, we used F4 generation of homozygotes to do viral infection. Transgenic materials were obtained by floral dipping with GV3101 contain PBA-Flag-4Myc-RAD51 in Col-0 or floral dipping with GV3101 contain pCambia1300-Myc-RAD51 in *atxr5 atxr6* as described ^58^.

### Plasmid constructs

Full-length coding DNA sequences (CDSs) of *RAD51* (AT5G20850), *CYCB1* (AT4G37490), *Rep* (encoded by CaLCuV) were cloned from CaLCuV-infected Col-0 and *HOP2* (AT1G13330) was cloned from U83877 (obtained from ABRC) into PENTER/D-TOPO (Thermo Fisher) vectors and confirmed by sequencing. The PCR was performed with *Thermococcus kodakaraenis* (KOD) DNA polymerase (Novagen).

Full-length CDSs of *BRCA1* (AT4G21070, stock G24692), *PARP1* (AT2G31320, G19185), *EMB1379* (AT5G21140, G63718), *TK1A* (AT3G07800, G10015), *TSO2* (AT3G27060, G11902), *XRI1* (AT5G48720, G63543), *HTA3* (AT1G54690, G13465), *HTA5* (AT1G08880, G13529), *PCNA1* (AT1G07370, G21468), RPA1A (AT2G06510, G61064), *PARG1* (AT2G31870, G16153), *WIP3* (AT4G10265, G60564) ligated with PENTER223 were obtained from ABRC and confirmed with enzyme digestion.

For Y2H constructs, full-length CDSs of *RAD51*, *CYCB1*, *Rep, HOP2*, *BRCA1*, *PARP1*, *EMB1379*, *TK1A*, *TSO2*, *XRI1*, *HTA3*, *HTA5*, *PCNA1*, *RPA1A*, *PARG1* and *WIP3* in PENTER-223 were cloned into pGADT7-DC and pGBKT7-DC by LR reaction. Plasmids confirmed by enzyme digestion were used for yeast transformation. pGADT7-ATXR5 and pGBKT7-ATXR5 were previously described^13^.

For transient expression constructs, full lengths CDSs of *RAD51*, *BRCA1*, *RPA1A, HOP2* were cloned into PBA-HA3-DC by LR reaction. Full length CDSs of *RAD51*, *CYCB1 and Rep* were cloned into PBA-Flag-4Myc-DC. Plasmids confirmed by enzyme digestion were transferred to GV3101 for transient expression in *N. benthamiana*.

For stable transgenic plants, PBA-Flag-4Myc-RAD51 was obtained as described above. To obtain pCambia1300-Myc-RAD51, pCambia1300-Myc-DC-Nluc was digested by PstI and dephosphorylated with Calf Intestinal (CIP) Alkaline Phosphatase. The missed destination cassette DC fragments missed was obtained by PCR of another normal DC cassette followed by digesttion by PstI. The resultant fragment was ligated into PstI/CIP treated pCambia1300-Myc-DC-Nluc to generate pCambia1300-Myc-DC. Full length CDS of RAD51 was transferred into pCambia1300-Myc-DC by LR reaction.

### CaLCuV infection assays

Plants were grown on MS plate under 8-h light/16-h dark condition for 10 days and then transferred to soil under 8-h light/16-h dark condition for around 14 days to reach eight-true - leaf developmental stage. Plants with different genotypes were infected by agroinfiltration of CaLCuV infective clones of PNSB1090 DNAA and PNSB1091 DNAB. The sympotoms of plants was monitored and evaluated daily since we began to observe the yellow mosaic and chlorosis in viral infected plants. To assess systemic infection, we harvested the eight newly emerged rosette leaves of CaLCuV-infected plants. To assess the plasmid transfection efficiency, we collected the whole plant except for cotyledons at 3, 6, 9, 13 and 16 dpi.

### Flow cytometry

Flow cytometry profiles were generated as described ^13^ with some modifications. We collected around 0.3 g rosette leaves from mock-treated plants with distinct genotypes at 14 dpi and finely chopped in 3 ml freshly made nuclear extraction buffer (45 mM Mgcl_2_, 30 mM sodium citrate, 20 mM pH 7.0 MOPS, 0.1% Triton X-100, 5 mM sodium metabisulfite, 5μl/ml 2-mercapto-ethonal, 100 μg/ml RNaseA) and filtered through 40 μm cell strainer (Sigma) to release the nuclei. Nuclei were stained by adding 150 μl 1mg/ml propidium iodide (Sigma) with gentle mixing by pipetting 10 times. Flow cytometry profiles were obtained from BD FORTESSA X-20 (College of Medicine Cell Analysis Facility, Texas A&M).

### Southern blot analyses

CaLCuV-infected plants were lysed in CTAB buffer (100 mM Tris-HCl pH 8.0, 20 mM EDTA, pH 8.0, 1.4 M Nacl, 3% cetyltrimethyl ammonium bromide, 2% β-mercaptoethonal) and then extracted with phenol:chlorophorm:isoamyl alcohol (25:24:1, pH 8.0) and precipitated with iso-propanol. Total DNA was treated with 50 μg/ml RNaseA and then purified with phenol:chlorophorm:isoamyl alcohol (25:24:1, pH 8.0). High quality DNA was obtained by precipitating with ethanol and sodium citrate. High quality DNA from plant was digested by EcoRI at 37 ℃ overnight. Digested DNA was transferred by capillarity to a Hybond-N membrane (GE Healthcare) and hybridized with ^32^P-labeled probe which is specific to CaLCuV DNAA. Probe for southern blot was amplified with primers listed in table S1 and then labeled using [α-^32^P] 2’-deoxycytidine 5’-triphosphate (dCTP) (PerkinElmer) with Klenow fragment (3’-5’ exo-; NEB). Hybridization signals were obtained using Typhoon FLA 7000 (GE Healthcare) and visualized in Photoshop.

### Western blot analyses

Western blot analyses were performed as previously described ^59^. Membranes were first incubated with antibodies against Myc (Sigma, C3956) or HA (Sigma, H9658) and then incubated with goat-developed anti-rabbit (GE Healthcare, NA934) or goat-developed anti-mouse immunoglobulin G (GE Healthcare, NA931). Membranes were developed with ECL+ and signals were obtained using ChemiDoc XRS+ and analyzed by Image Lab software (Bio-Rad).

### Strand Specific RNA sequencing library preparation

Total RNA was extracted from viral infected and mock treated plants at 16 dpi with TRIzol and then treated with TURBO Dnase (Thermo Fisher, AM2238). Messager RNA was enriched with Dynabeads™ Oligo(dT)_25_ (Invitrogen^TM^, 61005) according to the manufacturer’s manual. First strand synthesis was completed with Random primer (Invitrogen™, 48190011), RnaseIn (Invitrogen™, AM2696), SuperScript™ III Reverse Transcriptase (Invitrogen™, 18080044) after fragmentation. Second strand synthesis was conducted with dUTP mixure (20mM dUTP, 10mM dATP, dCTP, and dGTP), Rnase H (NEB, M0297) and DNA polymerase I (NEB, M0209). End repair was conducted with NEBNext Ultra^TM^ End repair/dA-Tailling Module (NEB, E7442) followed by the adapter ligation with ligation mix and adapter (Illumina, TruSeq Kits, 15026773) according to the manufacturer’s manual. Enriched mRNA from 1 μg total RNA was used as starting material and 15 cycles were used to amplify library.

### Chromatin Immunoprecipitation (ChIP) Assays

The ChIP assays were performed as previously described^3^ with some modifications. We collected eight newly emerged rosette leaves of mock-treated and viral-infected plants with different genotypes. Three grams of materials were crosslinked with 1% formaldehyde for 20 min by vacuum infiltration at 4 degree, and the reaction was stopped with a final concentration of 100 mM Glycine by vacuum infiltration for 10 min at 4 degree. Plants were rinsed 5 times with pre-chilled sterilized water, flash-frozen in liquid nitrogen, and thoroughly grounded with TissueLyser II (QIAGEN, 85300). The powder was suspended in pre-chilled 6 volume (1.5 g with 9 ml) nuclei isolation buffer (15 mM PIPES-KOH pH 6.8, 0.25 M sucrose, 0.9% Triton X-200, 5 mM MgCl2, 60 mM KCl, 15 mM NaCl, 1 mM CaCl2, 1 mM PMSF, 1 pellet/50 ml EDTA free Protease inhibitor [Roche] and 25 μM MG132) on ice for 6 min. The mixture was filtered through two-layer of Miracloth and centrifuged at 11000 g for 10min at 4 ℃ and carefully rinsed with 2 ml nuclei isolation buffer after discarding the supernatant. The white pellet was resuspended with 1 ml nuclei lysis buffer (40 mM Tris-HCl pH 8.0, 150 mM NaCl, 5 mM EDTA pH 8.0, 0.2% SDS, 0.1% Sodium Deoxycholate, 0.6% Triton X-100, 1 mM PMSF, 1 pellet/50 ml EDTA free Protease inhibitor [Roche] and 25 μM MG132) on ice for 6 min and then sonicated with 30 s ON and 60 s OFF for 16 cycles with Bioruptor Pico sonication device (Diagenode SA, B01060010) at 4 ℃. Supernatant was collected after centrifuge for 10 min at 15000 rpm at 4 ℃ and diluted with the same volume nuclei dilution buffer (40 mM Tris-HCl pH 8.0, 150 mM NaCl, 5 mM EDTA. 0.2% Triton X-100, 1 mM PMSF, 1% Glycerol, 1 pellet/50 ml EDTA free Protease inhibitor [Roche] and 25 μM MG132). 100 μl diluted mixture was used as input, then the rest was split into two tubes with addition of 30 μl prewashed Protein A Agarose beads (Sigma, 11134515001), 3 μg anti-RAD51 (PHYTOAB, PHY1804A) and 3 μg anti-RPA1A (PHYTOAB, PHY1813S), and incubated at 4 degree with mild rotation for 6 hours. The beads-protein-DNA complex was washed as previously described ^3^. Elution was repeated twice at 1200 rpm, 65 ℃ with 250 μl elution buffer (0.1 M NaHCO3, 1% SDS) for 30 min. De-crosslinking was performed with 100 NaCl at 65 ℃ for 12 hour, followed by Rnase A treatment and Protease K treatment. Purification was performed with phenol:chlorophorm:isoamyl alcohol (25:24:1, pH 8.0), precipitation of DNA was performed with 1/10 volume of 3 M sodium citrate (pH 5.2), 2.5 volume of Ethyl alcohol and 1μl GlycoBlue (Invitrogen ^TM^, AM9516) and DNA was dissolved in 1×TE buffer (10 mM Tris-HCl pH 8.0, 1 mM EDTA pH 8.0).

### Quantitative PCR

Relative amount of viral DNA A in viral-infected plants and endogenous rDNA of *Arabidopsis* were examined by q-PCR. High quality DNA was obtained as described in southern blot analyses. *Ubiquitin 10* served as an internal control for normalization. The enrichment levels of RAD51 and RPA1A on specific loci on viral genome and *Arabidopsis* after ChIP assays were also accessed by q-PCR. The q-PCR assays were performed with 10 μl system contain 5 μl SYBR Green Master Mix in 384-well plate. PCR conditions, signal detection and quantification were performed as previously described ^3^.

### ChIP sequencing library preparation

Half of ChIP-enriched DNA or one sixth of input DNA were used as starting materials for DNA end repair process for library construction with NEBNext Ultra^TM^ End repair/dA-Tailling Module (NEB, E7442). Then the products were ligated to adapters (NEBNEXT Multiplex Oligos for Illumina, E7335, E7500 and E7710) with Blunt/TA Ligase Master Mix (NEB, M0367) according to the manufacturer’s manual. For the library preparation, 9 cycles were used to amplify products from input DNA, 13 cycles were used to amplify RAD51 ChIP-enriched DNA and 16 cycles were sued to amplify RPA1A ChIP-enriched DNA.

### Illumina Sequencing and analysis

Libraries for strand specific RNA sequencing were sequenced on the Illumina Hiseq 2500 platform with pair-end 50 bp read length (Novogene). Adapters trimming, mapping and counting process were performed as previously described^13^. Normalization and differential expressed analysis were calculated by DESeq2 (ver.3.15)^60^. The visualization for selected loci was shown in Intergrative Genomics Viewer (IGV)^61^.

Heat-map clustering was performed based on Pearson distance correlation. To obtain the comprehensive DEG lists which contain both mock-treated and viral infected samples, We first compared mock-treated *atxr5 atxr6* to mock-treated Col-0 using Col-0 as a reference, and then compared the other three triple mutants to *atxr5 atxr6* using *atxr5 atxr6* as a reference. The same analysis was also performed on viral infected samples. Moreover, we also compared the viral infected Col-0 to mock-treated Col-0 using as a reference, then compared the other viral infected genotypes to their mock-treated samples. The DEGs were selected using the cut off (Fold change>2 and *p*<0.05), but the genes with low expression (max reads>10) were filtered. Pool of DEGs from the three groups mentioned above were the comprehensive DEG lists to perform heatmap clustering analysis.

Libraries for ChIP-seq were sequenced on the Illumina Hiseq 2500 platform with pair-end 150 bp read length (Novogene). Adapter trimming and mapping processes were performed as previously described ^13^. The leftover reads later were mapped by bowtie2 (ver. 2.4.4) ^62^ with perfect matches using Arabidopsis genome TAIR10 (http://www.arabidopsis.org/) as a reference genome. Reads uniquely mapped to genome and reads mapped to multiple locations in the genome were separately extracted by Samtools (ver. 1.15.1)^63^ for distinct downstream peak calling analyses. MACS2 (ver. 2.2.5)^64^ was used for ChIP-seq peak calling using default parameters, both narrowPeak and broadPeak were called for each sample. We used the reads mapping to genome which contain sequence mapped to multiple locations in the genome to perform the ChIP signal profile, peak number, peak reads percentage analysis and comparative analysis with DESeq2 (ver.3.15)^60^. Peaks with *P-*value < 0.05 were selected for the following analysis, narrowPeak and broadPeak were merged by bedtools^65^ (ver. 2.29.2).

Annotation of peaks was conducted using bedtools. with TAIR10 gtf file as a reference. Peaks from biological replicates for the same genotype were merged and classified into different categories to calculate the relative peak number percentage. The peaks belonging to distinct categories were first extracted with Samtools^63^, and then reads over peaks from distinct categories were counted with Subread feature Counts (ver. 2.0.0)^66^. Total reads of peaks from specific categories were divided by total reads of peaks from all categories to generate peak reads percentages for individual specific categories.

All downstream statistical analyses and plotted graphs were generated by R (ver. 4.0.2) and ggplot2^67^. All ChIP-seq profiles for RAD51, RPA1A, and H3K27me1 were drawn by deeptools (ver. 3.7.4)^68^ using the default parameters. DESeq2 (ver.3.15)^60^ with default parameters (*P-*value < 0.05) was implemented to output differential peaks between different groups of ChIP-seq samples. All Violin plots were plotted by ggviolin package. All the plots and profiles for Chip-seq were drawn by R. The visualization for selected loci was shown in Intergrative Genomics Viewer (IGV)^61^.

### Yeast Two-Hybrid (Y2H) Assays

Y2H were performed using the Gold Yeast Two-Hybrid System according to manufacturer’s manual (Clontech). CDSs from different genes with combination of pGADT7 and pGBKT7 were co-transformed into the yeast strain AH109. The yeast transformants were simultaneously plated on medium minus Leu, Trp, His, Ade to screen the interactions and medium minus Leu, Trp as controls under 28 ℃.

### Co-immunoprecipitation (Co-IP) Assays

GV3101 with tested constructs were co-transformed into *N. benthamiana* leaves by agroinfiltration. Materials were collected at two days post transformation, grounded in liquid nitrogen and stored at −80℃. Total proteins were extracted from 0.35 g powder in 1.0 ml IP buffer and then centrifuged twice at 15,000 g for 15 min at 4 ℃. IP buffer for HA-RPA1A and FM-Rep, HA-Rep and FM-RAD51, HA-RPA1A and FM-RAD51 was 40 mM Tris-HCl pH 8.0, 150 mM NaCl, 5 mM MgCl_2_, 1% Glycerol, 0.6% Triton X-100, 1 mM PMSF, 1 pellet/50 ml EDTA free Protease inhibitor [Roche] and 25 μM MG132. IP buffer for HA-BRCA1 and FM-RAD51, HA-HOP2 and FM-RAD51, HA-RAD51 and FM-CYCB1 was 40 mM Tris-HCl pH 7.0, 150 mM NaCl, 5 mM MgCl_2_, 1% Glycerol, 0.6% Triton X-100, 1 mM PMSF, 1 pellet/25 ml EDTA free Protease inhibitor [Roche] and 50 μM MG132. Anti-HA-agarose beads or anti-FLAG-M2 magnetic beads were added to total protein extracts and then incubated at 4 ℃ for 2 h. Turbo DNases (10 U/ml) was added to IP buffer prior to incubation to conduct DNase treatment. The beads were washed four times at 4 ℃ for 5 min with IP buffer after incubation, then boiled with SDS loading buffer at 95 ℃. Western blot analyses were carried out as described above with anti-Myc (Sigma, C3956) or anti-HA (Sigma, H9658) to detect IP products and co-precipitation levels of their partners.

### Quantification and Statistical analysis

The images of southern blot were quantified with Gel-Pro Analyzer (Media Cybernetics). For RNA-seq and ChIP-seq, an DESeq2 (version 3.15)^60^package was used to normalize gene expression levels with trimmed mean of M-values according to the false discovery rate. The significance cutoff for RNA-seq was Log2 FC>1 or Log2 FC<-1, *P*-value< 0.05, and for ChIP seq, the cutoff was Log2 FC>0.5 or Log2 FC<-0.5, *P*-value< 0.05,

For q-PCR, the data were presented as means of at least three replicates ±SD. For Fig. 2G, 3G, 5I, 6G and S7D, the relative amount of viral DNA was initially normalized to that of *UBQ10*, and then to WT (Col-0) where the ratio was arbitrarily assigned a value of 1 with ±SD (*n* = 3) biologically independent replicates. For Fig. S12C, the amount of rDNA was normalized to that of *UBQ10*.

### Graph drawing

Go enrichment analysis were performed using Metascape^69^. PPI network prediction was performed in STRING^70^ and polished in Cytoscape^71^. Graphs with dot plots (individual data points) were drawn using GraphPad Prism 9. Model was drawn using BioRender.

### Data availability

RNA-seq related data were mined from GSE77735^12^.H3K9me2 and H3K27me1 related data were mined from GSE111814^13^. H3K27Ac related data were mined from GSE146126^10^ and H3K4me3 related data were mined from GSE166897^14^. RAD51 and RPA related data were mined from GSE143582^36^. The data generated during this study will be deposited in GEO once the manuscript is accepted.

## Supplementary Information

**Extended Data Fig.1 Loss of function mutants of *ATXR5 and ATXR6*, different from other epigenetic mutants, displayed viral resistance phenotype.**

**Extended Data Fig.2 Transfection efficiency was comparable between Col-0 and *atxr5 atxr6*.**

**Extended Data Fig.3 H3.1 single and quadruple mutants and *fas-2*, unlike *atxr5 atxr6*, were hyper-susceptible to viral infection.**

**Extended Data Fig.4 The molecular phenotype of TE reactivation is uncoupled with the viral resistance phenotype in *atxr5 atxr6*.**

**Extended Data Fig.5 Highly reproducible datasets High quality of RNA-seq.**

**Extended Data Fig.6 TWAS analysis identified the genes that are highly correlated with viral resistance of *atxr5 atxr6*.**

**Extended Data Fig.7 Y2H screening of HRR proteins identified a few Rep-interacting proteins.**

**Extended Data Fig.8 Quality analysis of ChIP-seq for RAD51 and RPA1A.**

**Extended Data Fig.9 Distributions of RAD51-bound peak over euchromatin and heterochromatin regions.**

**Extended Data Fig.10 ChIP signal of RAD51 and RPA1A in mock-treated Col-0 and *atxr5 atxr6*.**

**Extended Data Fig.11 Gene Ontology (GO) analysis of RAD51 and RPA1A enriched PCGs.**

**Extended Data Fig.12 The change of RAD51 and RP1A signal induced by viral infection in Col-0 and *atxr5 atxr6*.**

**Extended Data Fig.13. Viral infection enhanced the RAD51 signal over PCGs related to defense response.**

**Extended Data Fig.14 Deletions of HOP2, BRCA1 and CYCB1 do not affect the viral DNA replication.**

**Supplementary Excel 1.** Gene lists used in Fig. 2, a and b, and Fig. 2d, and Fig. 4h, and Fig. 5c, and Extended Data Fig. 6d, and Extended Data Fig. 13, a, b and d.

